# Systems Modeling Reveals Shared Metabolic Dysregulation and Potential Treatments in ME/CFS and Long COVID

**DOI:** 10.1101/2024.06.17.599450

**Authors:** Gong-Hua Li, Feifei Han, Efthymios Kalafatis, Qing-Peng Kong, Wenzhong Xiao

## Abstract

Myalgic Encephalomyelitis/Chronic Fatigue Syndrome (ME/CFS) and Long COVID are complex, multisystem conditions that pose significant challenges in healthcare. Accumulated research evidence suggests that ME/CFS and Long COVID exhibit overlapping metabolic symptoms, indicating potential shared metabolic dysfunctions. This study aims to systematically explore shared metabolic disturbances in the muscle tissue of patients. Utilizing genome-wide metabolic modeling, we identified key metabolic irregularities in the muscle of patients with ME/CFS, notably the downregulation of the alanine and aspartate metabolism pathway and the arginine and proline metabolism pathway. Further, *in silico* knockout analyses suggested that supplementation with aspartate (ASP) or asparagine (ASN) could potentially ameliorate these metabolic deficiencies. In addition, assessments of metabolomic levels in Long COVID patients also showed the significant downregulation of ASP during post-exertional malaise (PEM) in both muscle and blood. Consequently, we propose that a combination of l-ornithine and l-aspartate (LOLA) is a potential candidate to alleviate metabolic symptoms in ME/CFS and Long COVID for future clinical trials.

## Introduction

The emergence of Myalgic Encephalomyelitis/Chronic Fatigue Syndrome (ME/CFS) and Long COVID as prominent health crises has underscored the need for a deeper understanding of these complex, multisystemic conditions. ME/CFS, a debilitating chronic condition that has been historically under-studied, is characterized by persistent and unexplained fatigue, post-exertional malaise (PEM), orthostatic intolerance, unrefreshing sleep, brain fog, muscle pain, and other symptoms^1-3^. While Long COVID represents a spectrum of symptoms severely impacting the quality of life of patients, persisting months beyond the acute phase of SARS-CoV-2 infection^4,5^, most of the symptoms are similar to those of ME/CFS, with few exceptions^6-8^.

The similar clinical presentations of ME/CFS and Long COVID have prompted researchers to hypothesize and explore shared underlying mechanisms, particularly within metabolic functions. Metabolic dysfunction has been increasingly recognized as a potential contributor to the symptomatology of these conditions, with disturbances noted in energy metabolism, amino acid and lipid profiles, and mitochondrial function^9-14^. Skeletal muscle, together with the liver and the brain, consumes the most energy at rest and dramatically increase its energy consumption during physical exertion^15^. Investigations of muscle tissues are likely critical in understanding the metabolic mechanism underlying the key symptoms of these diseases and identifying candidates for potential treatments.

In this study, we investigate the shared metabolic alterations in the muscle of ME/CFS and Long COVID patients by using genome-wide precision metabolic modeling (GPMM)^16^ and metabolomic data analysis (**Figure 1**). We identified that the most significant metabolic change is the downregulation of alanine and aspartate metabolism. We also propose the combination of l-ornithine and l-aspartate (LOLA) as a potential therapeutic candidate to replenish these deficient metabolic pathways.

**Figure 1:**
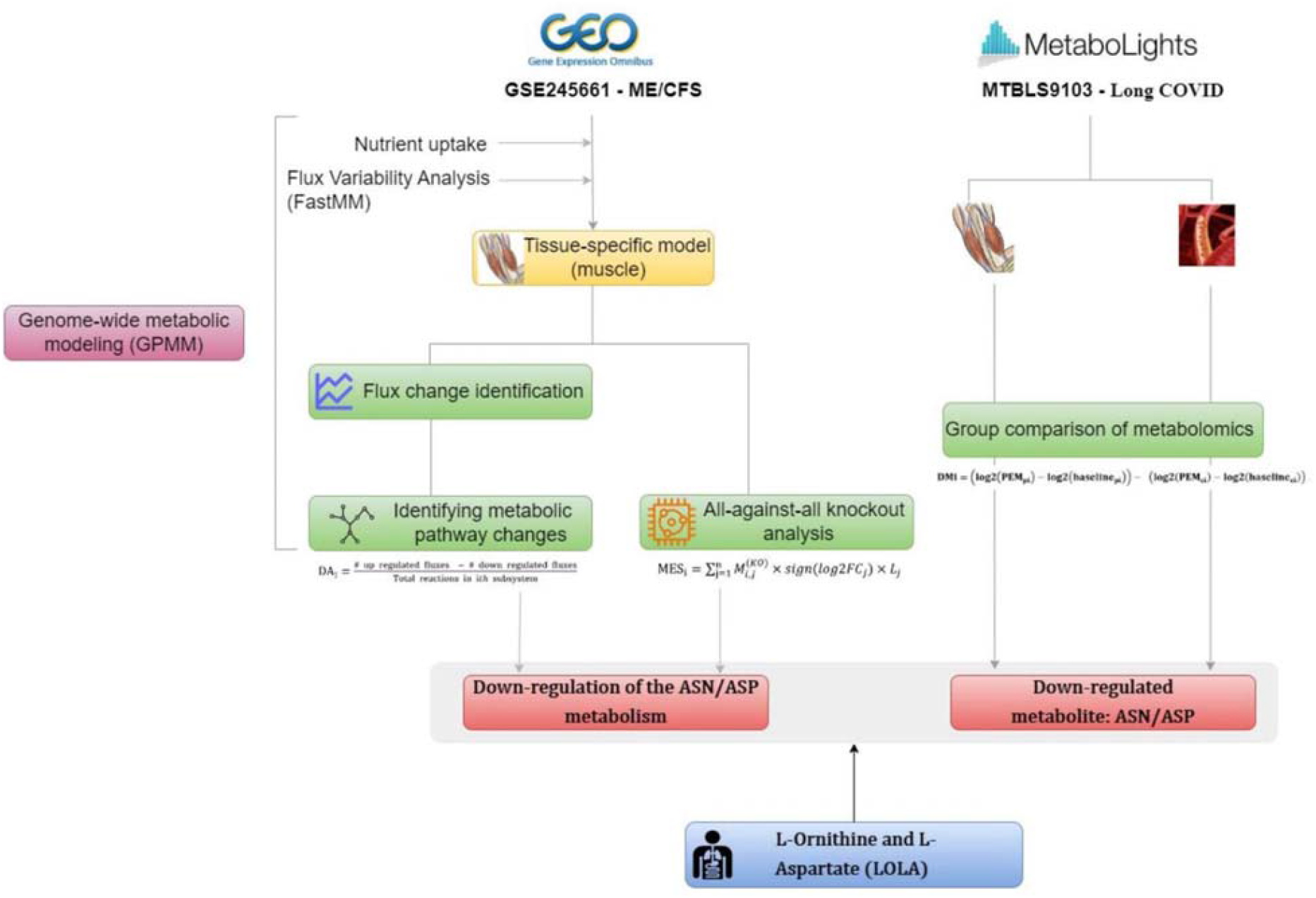
The workflow of Genome-Wide metabolic modeling (GPMM) and Metabolomics leading to target identification for ME/CFS and Long COVID. The left panel showed GPMM process using transcriptomic data of muscle tissues from an ME/CFS study, GSE245661. The right panel showed the data analytics process of metabolomics data from a Long COVID study, MTBLS9103. Both processes identified ASN/ASP metabolism as the most down-regulated pathway, resulting in the finding of LOLA as a potential treatment candidate for both diseases.

## Results

### Metabolic Modeling Reveals Altered Metabolism in the Muscle of ME/CFS Patients

To investigate metabolic changes in ME/CFS, we conducted metabolic modeling using muscle samples from ME/CFS patients and controls from the NIH deep phenotyping of ME/CFS study^17^. The metabolic model of the muscle tissue involved 2,841 metabolic reactions, 2,248 enzymes, and 1,519 metabolites.

Comparing the flux of each metabolic reaction between patients and controls identified 65 reactions showing significant up-regulation and 50 reactions showing significant down-regulation in the muscle of patients (**Figure 2A**, p<0.05, |log2fc| > 0.2) (**Supplemental Data 1**). Notably, among the downregulated fluxes, four (ASPNATm, ASPTAm, NACASPAH, ASPTA) were related to alanine and aspartate metabolism, and two (4HGLSDm and PHCHGSm) were related to arginine and proline metabolism. Among the upregulated fluxes, seven (G6PDH2r, PGL, RPE, RPI, TKT1, TKT2, GND) were related to the pentose phosphate pathway.

**Figure 2:**
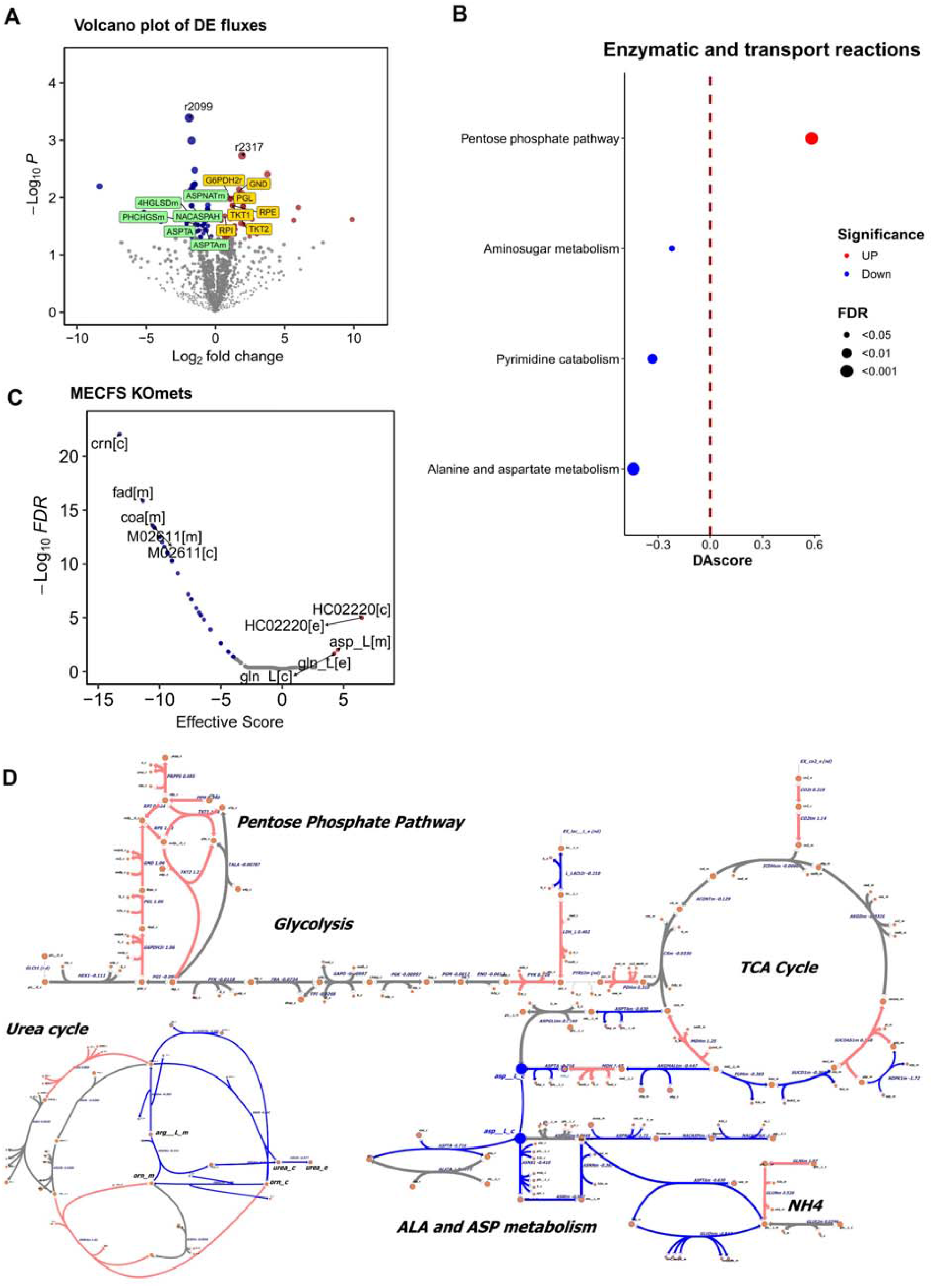
Metabolic modeling of muscle samples of ME/CFS patients. (A) volcano plot of differential changed metabolic fluxes between ME/CFS patients and controls (p-value<0.05, |log2fc| > 0.2). Red, blue and gray points represent significantly increased, decreased, and non-significant pathways, respectively. Four fluxes related to alanine and aspartate metabolism (ASPNATm, ASPTAm, NACASPAH, ASPTA) and two fluxes related to arginine and proline metabolism (4HGLSDm and PHCHGSm) were downregulated and colored by light green. Seven fluxes related to the pentose phosphate pathway (G6PDH2r, PGL, RPE, RPI, TKT1, TKT2, GND) were significantly upregulated and colored by light yellow. Note that r2099 is a transport reaction (Tcdb:2.A.1.13.1) mediated by the SLC16A1 gene, and r2317 is also a transport reaction (Tcdb:2.A.60.1.2) mediated by the SLCO2A1 gene. (B) Pathway analysis differential changed metabolic fluxes between ME/CFS patients and controls (FDR<0.05 and |DAscroe| > 0.2). Red, blue and gray points represent significantly up-regulated, down-regulated, and non-significant pathways, respectively. (C) Metabolite knockout analysis in ME/CFS. Effective scores greater than 0 indicate agonist metabolites, while scores less than 0 indicate antagonist metabolites. Administering an agonist metabolite, e.g. ASP, could potentially rescue the changes observed in patients. (D) Metabolic map of representative pathways metabolic fluxes. The down-regulated reactions are enriched on Alanine and aspartate metabolism in patients. In addition, the down-regulated reactions include reduced urea secretion and a potential increase in NH4 secretion. On the other hand, the up-regulated reactions are enriched on the pentose phosphate pathway.

Pathway enrichment analysis further revealed that several metabolic pathways were affected. Specifically, pathways of alanine and aspartate metabolism, pyrimidine catabolism, aminosugar metabolism were significantly down-regulated, and conversely, the pentose phosphate pathway was significantly up-regulated (FDR < 0.05, |DAscore| > 0.2). Notably, the most prominently down-regulated pathway was alanine and aspartate metabolism in the muscle of ME/CFS patients (**Figure 2B**).

Next we examined the metabolic network of associated with glucose and amino acid metabolism. Intriguingly, most of the reactions in alanine and aspartate metabolism are down-regulated (log2FC < -0.2) in the muscle sample of ME/CFS patients (**Figure 2D**). Additional flux changes (e.g. urea_c and urea_e in the urea cycle) indicated reduced urea secretion and a potential increase in NH4 secretion (labeled as NH4). On the other hand, the up-regulated reactions were clearly enriched on the pentose phosphate pathway.

We further investigated whether increasing the metabolites on Alanine and Aspartate metabolism pathway can potentially rescue the metabolic changes seen in patients toward the metabolic state of the controls. Our all-against-all knockout analysis highlighted that Asparagine (ASN) and Aspartate (ASP) are also among the top agonist metabolites (**Figure 2C**) (**Supplemental Data S2**), suggesting that administering these two amino acids could potentially rescue the metabolic changes observed in ME/CFS patients. This knockout result aligned with the modeling findings and supported the notion of dysfunction in alanine and aspartate metabolism among ME/CFS patients^10^.

### Metabolomics Measurements of Long COVID Reveals Down-Regulated Asparagine (ASN) during PEM in Muscle and the Blood

Given the similarity in symptoms between long COVID and ME/CFS, we analyzed the metabolic profile of the muscle in Long COVID patients. Since PEM is a cardinal symptom of ME/CFS and common in Long COVID, we performed pairwise comparisons between pre- and post-exercise samples from Long COVID patients and healthy controls using data from a study of muscle tissue during post-exertional malaise in Long COVID^18^(**Supplementary Data S3**). As shown in **Figure 3A**, significant changes were noted in muscle tissue, particularly the significantly lower level of asparagine and dihydroxyacetone-P during PEM. Similarly, in blood samples, Asparagine also ranked highest among down-regulated metabolites following PEM in Long COVID patients (**Figure 3B**).

**Figure 3:**
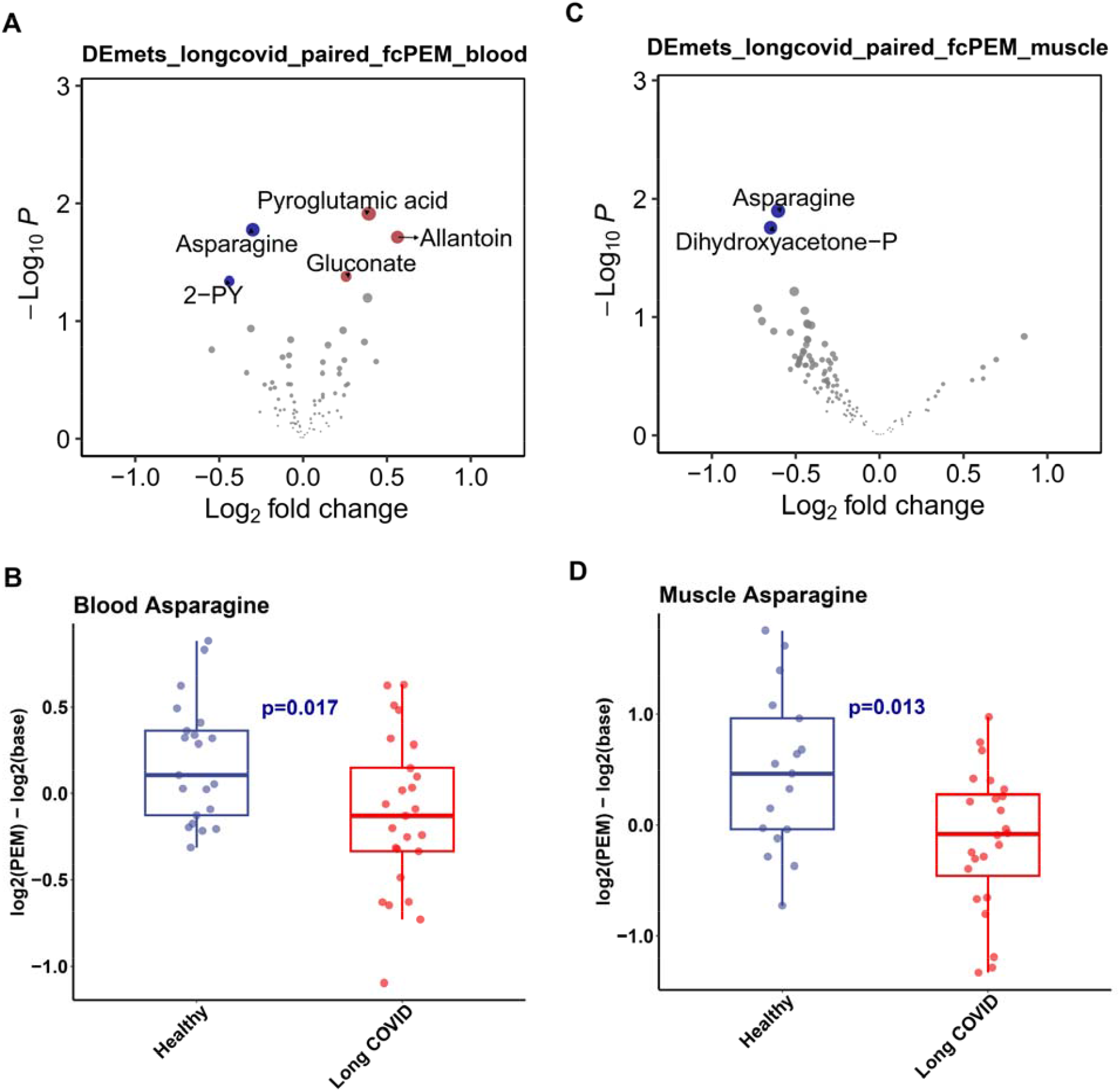
Pair-wise changes of metabolite levels in Long COVID after post-exertional malaise (PEM). (A). Volcano plot of pair-wise metabolic changes in the muscle of Long COVID patients after PEM. (B). Boxplot of changes of asparagine level in the blood between Long COVID and healthy controls. (C). Volcano plot of pair-wise metabolic changes in long-covid patient blood after PEM.(D). Boxplot of changes of asparagine in the muscle between Long COVID and Healthy controls.

Collectively, these consistent findings from the ME/CFS and the Long COVID studies highlighted the significant down-regulation of the ASN/ASP metabolism in the muscle tissue of both conditions.

### L-Ornithine and L-Aspartate (LOLA) as a Potential Treatment Candidate for ME/CFS and Long COVID

Next, we propose a candidate of potential treatment targeting these specific metabolic pathways for both ME/CFS and Long COVID. We suggest that LOLA may offer therapeutic benefits for individuals with these conditions for several reasons (**Figure 4**):

**Figure 4:**
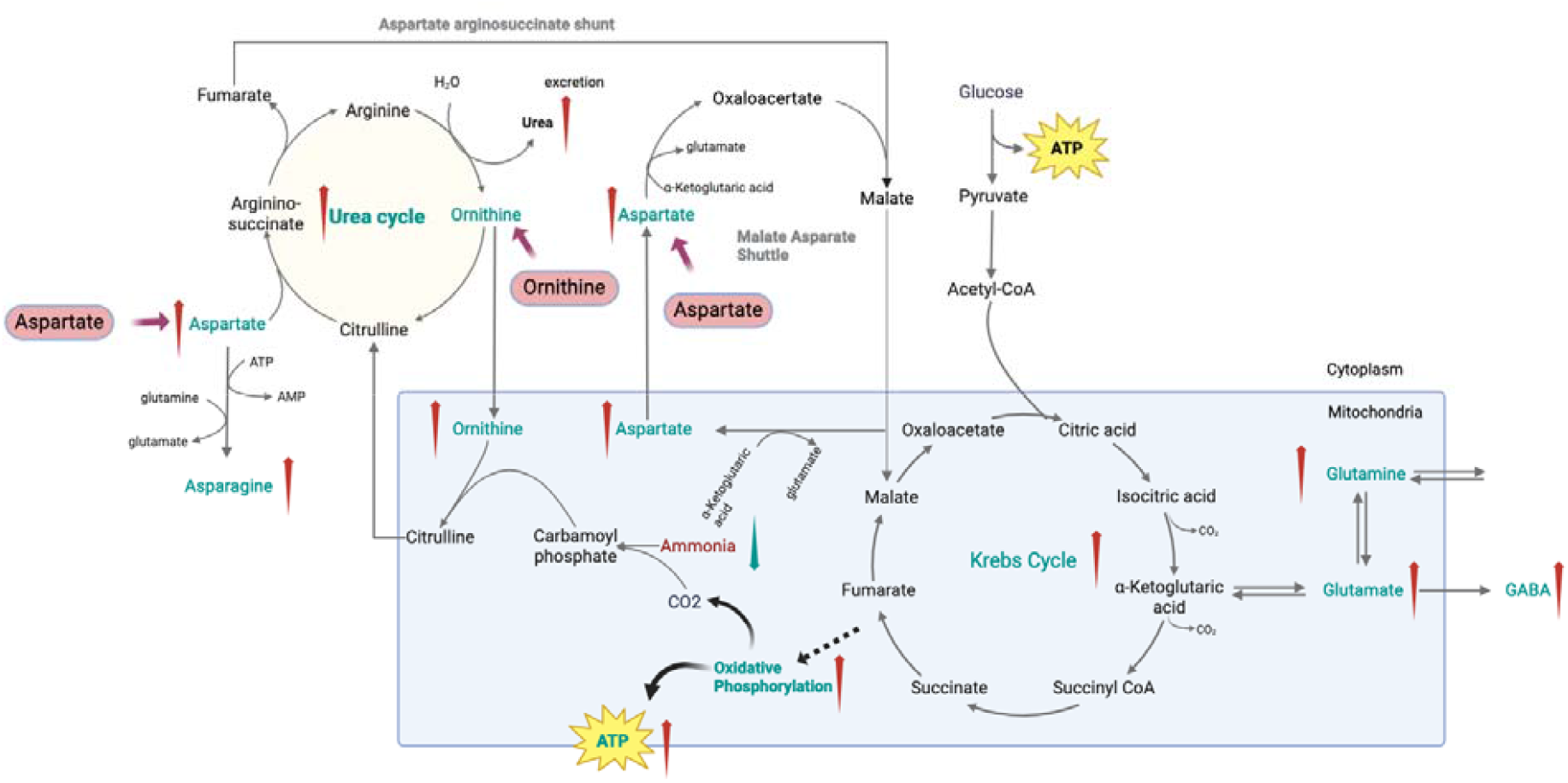
L-Ornithine and L-Aspartate (LOLA) as a Potential Treatment Candidate for ME/CFS and Long COVID. L-Aspartate potentially can help counteract the observed decrease in Asparagine/Aspartate in both ME/CFS and Long COVID. L-ornithine, as the metabolic product of arginine, potentially can help restoring the balance of the down-regulated pathway of arginine and proline metabolism in ME/CFS. Additionally, the combined use of these amino acids could enhance urea cycle efficiency, which is critical in removing ammonia and reducing fatigue symptoms commonly reported in these conditions. Besides emerging evidence suggests LOLA can improve mitochondrial function, thereby potentially enhancing energy metabolism in patients with ME/CFS and Long COVID.

1. L-Aspartate aligns with the commonly observed down-regulation of ASN/ASP in both ME/CFS and Long COVID^19^, suggesting it could help counteract this deficiency.
2. As the metabolic product of arginine, l-ornithine corresponds to the down-regulated pathway of arginine and proline metabolism in ME/CFS, potentially restoring balance in this pathway^20^.
3. Additionally, the combined use of these amino acids in LOLA could enhance the efficiency of the urea cycle ^21^, which is critical in removing ammonia and reducing fatigue symptoms commonly reported in these conditions^9^.
4. Emerging evidence suggests that supplementation with LOLA can improve mitochondrial function^22,23^, thereby potentially enhancing energy metabolism, which is often impaired in patients with ME/CFS and Long COVID^24^.

## Discussion

This study provides an analysis of the metabolic disruptions found in both ME/CFS and Long COVID, offering new insights into their pathophysiology and highlighting potential treatment avenues. Our findings reveal significant metabolic commonalities between these conditions, particularly in the down-regulation of amino acid metabolism pathways such as ASN/ASP and arginine/proline in the muscles. These findings not only help us understand better the systemic impact of these conditions but also highlight potential targets for therapeutic intervention.

The consistent down-regulation of specific metabolic pathways across both indications suggests a fundamental disruption in amino acid metabolism and energy metabolism, which could be contributing to the severity and persistence of patients’ symptoms^25^. In addition, Asparagine provides key sites for N-linked glycosylation, which is required for proper protein folding in the endoplasmic reticulum (ER)^26^, and otherwise may induce ER stress, which likely takes place in ME/CFS^27^.

Here, we propose that L-ornithine and L-aspartate (LOLA) might hold the potential of intervening these metabolic pathways. L-aspartate, for instance, could directly replenish the decreased aspartate pool, while L-ornithine might work by restoring the urea cycle pathway, thereby improving the overall metabolic balance and reducing symptoms such as fatigue and cognitive dysfunction^28,29^.

Moreover, the role of mitochondrial dysfunction in these conditions cannot be ignored either. Recent studies have shown that mitochondrial dysregulation is a key factor in the pathogenesis of chronic diseases, including those characterized by post-exertional malaise, a hallmark of both ME/CFS and Long COVID^24^. By enhancing mitochondrial function and energy production, LOLA could potentially mitigate some of the core symptoms of these conditions.

Furthermore, ammonia dysregulation has also been implicated in fatigue and cognitive dysfunction observed in both ME/CFS ^30-32^ and Long COVID patients^33,34^. Hyperammonemia is known to occur after intense or exhausting exercise and in pathological liver disorders^35^, and it can affect energy production^36^ and potentially induce neuroinflammation^37^. LOLA has been studied clinically in these conditions to facilitate ammonia detoxification in the liver by replenishing intermediates of the urea cycle^21,38^. It may help alleviate the metabolic symptoms associated with ammonia dysregulation in ME/CFS and Long COVID.

This study highlights the importance of investigating skeletal muscle, a major metabolic organ system, to better understand the metabolic dysfunctions in complex multisystem conditions. Further research of major metabolic tissues could pave the road for personalized medicine strategies in treating ME/CFS and Long COVID, where treatments are tailored based on specific metabolic profiles of patients.

Clinical trials are essential for evaluating the long-term efficacy and safety of LOLA in these conditions, and longitudinal tracking of metabolic alterations and key symptoms in patients over time will be invaluable in validating and further elucidating its potential effects. Notably, this study focuses on analyses of skeletal muscles of patients and the potential impact of LOLA on the metabolic system. Exploring the interactions between the metabolic and other physiological systems, such as the immune, endocrine, and neurological systems, is crucial to developing a deeper understanding of these complex conditions and discovering better treatments.

In addition to LOLA, sulfochenodeoxycholate (HC02220), a sulfoconjugated chenodeoxycholic acid (CDCA) was also identified as an agonist (**Figure 2C**). CDCA is a precursor in the formation of taurochenodeoxycholic acid (TUDCA), which potentially improves insulin sensitivity and supports mitochondrial function in the skeletal muscle^39^. Further study is required to further investigate these metabolites in ME/CFS and long COVID.

In conclusion, by systematically exploring the metabolic mechanism in the muscles of patients, our study contributes to understanding of the underlying mechanism of the complex conditions and candidates for potential treatments. Future clinical trials of LOLA will evaluate its safety and efficacy in alleviating metabolic symptoms in ME/CFS and Long COVID patients.

## Resource availability

### Lead contact

Further information and requests for resources should be directed to and will be fulfilled by the lead contact, Wenzhong Xiao.

### Materials availability

This study did not generate new unique reagents.

## Data and code availability

Data used in this research are all publicly available. Accession numbers are listed in the key resources table.

Metabolic modeling, flux analysis and visualization, the metabolic pathway analysis, all-against-all knock out analysis, and key metabolite identification were all performed by rGPMM (https://github.com/GonghuaLi/rGPMM) with the version 1.0.0 on the metabolic models Recon3, version 1 using the CPLEX solver. Individual metabolic flux can be queried online at http://bigg.ucsd.edu/models/Recon3D/reactions. The code for this manuscript can be available at: https://github.com/GonghuaLi/Code_for_publications/tree/master/MECFS_LOLA.

Any additional information required to reanalyze the data reported in this paper is available from the lead contact upon request.

## STAR★Methods

### Key resources table

| REAGENT or RESOURCE | SOURCE | IDENTIFIER |
| --- | --- | --- |
| **Deposited data** |  |  |
| Transcriptome dataset of muscle tissues of ME/CFS and controls | GEO | GSE245661 |
| Metabolomics dataset of the muscle and blood samples of Long COVID | MetaboLights | MTBLS9103 |
| **Software and algorithms** |  |  |
| rGPMM | https://github.com/GonghuaLi/rGPMM | v1.0.0(need request from lead contact) |

### Method details

#### Dataset collection

The transcriptome dataset of muscle tissues of ME/CFS and controls was accessed from GEO (GSE245661)^17^. This was part of the NIH deep phenotyping of post-infectious ME/CFS study^17^, and comprises of RNA sequencing data of 13 ME/CFS patients and 12 healthy controls. The metabolomics dataset of the muscle and blood samples of Long COVID was obtained from MetaboLights (MTBLS9103)^18^. The dataset we analyzed includes metabolomics measurements of skeletal muscle biopsies of 46 long COVID patients taken before and 1 day after exercise testing along with the metabolomics measurements of the blood samples. Detailed protocols of muscle biopsies, exercise testing, and genomic and metabolomic measurements are in the respective publications^17,18^.

#### Genome wide precision modeling of metabolic fluxes in the muscle of patients and controls

##### Metabolic modeling

We used our recently developed genome-wide precision metabolic modeling method, GPMM, to perform the genome-wide metabolic modeling^16^. Briefly, GPMM integrates protein abundance estimates from gene expression data with enzymatic kinetic parameters, and uses these as upper bounds in a generic human metabolic model based on Michaelis–Menten kinetics. Nutrient uptake fluxes for cell lines were referenced from existing literature^40^, and the lower bounds for other exchanges were set as zero. The metabolic model in Recon3 (version 1) ^41^ was utilized as the initial general model of human metabolism. We conducted Flux Variability Analysis (FVA) using the FastMM algorithm to construct tissue-specific models for each sample^42^.

##### Identification of significantly changed fluxes

For case-control design, the flux changes between patients and controls were calculated using the Limma package ^43^ based on the resulting values of fluxes from the metabolic modeling. Differentially expressed (DE) fluxes with Pvalue <0.05 were identified as significantly changed.

##### Identification of significantly changed metabolic pathways

To ascertain changes in metabolic pathways, we calculated the Differential Abundance Score (DA score) employing a method previously described in the literature ^44^. For each metabolic pathway (i), the DA score (DA_i_) is computed as follows:

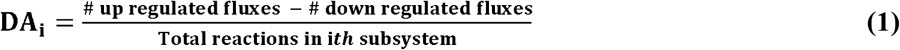

The statistical significance (p-values) of the Differential Abundance (DA) scores was determined using a “bootstrap without replacement” method, as detailed in our previous study^16^.

##### All-against-all knockout analysis

We performed an all-against-all metabolite knockout analysis using the FastMM^42^ algorithm to generate a comprehensive metabolite knockout matrix (M^(KO)^). In this matrix, rows represent metabolites, while columns correspond to reactions. The Metabolite Effective Score (MES) was calculated in a manner analogous to gene knockout analysis, as follows:

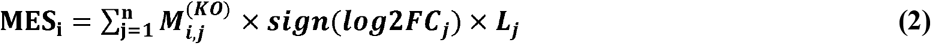

Here, 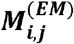 denotes the effect score of the *ith* metabolite on the *jth* reaction. The significance (p-value) of each MES was determined using a norm-background method, employing the pnorm function in R. FDR adjustments were made using the p.adjust function with the ‘FDR’ method in R. An agonist, identified as its MES value greater than 0 and an FDR less than 0.05, is a metabolite the knockout of which exacerbates the flux changes seen in patients, suggesting that administering the metabolite could potentially rescue the changes observed in patients. On the other hand, an antagonist, identified as its MES value lower than 0 and an FDR less than 0.05, is a metabolite the knockout of which reduces the flux changes seen in patients, suggesting that reducing the metabolite could potentially rescue the changes observed in patients.

#### Metabolomics data analysis

In MTBLS9103, 116 metabolites in skeletal muscle and 83 metabolites in venous blood were annotated^18^. For both tissues, we calculated Differential Metabolite (DM) using equation (3) to quantitate the difference of metabolite concentration change after PEM between Long COVID patients and controls. The significance was calculated by employing the eBayes method using the limma package in R^45^.

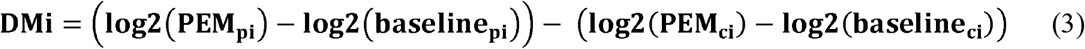

## Acknowledgments

This work was supported by grants from Open Medicine Foundation (WX). Yunnan Ten Thousand Talents Plan Young & Elite Talents Project (G.-H.L.), Yunnan Fundamental Research Projects (202101AS070058). We would also like to express our sincere gratitude to the patients who provided valuable feedbacks to us on the results of this study.

## Author contributions

WZX and GHL conceived and designed the study. WZX, GHL and QPK supervised the project. GHL designed and developed the computational method. GHL, FFH and WZX collected the datasets. GHL and FFH performed the data analysis and conducted the statistical analysis. GHL, FFH, and WZX drafted the manuscript. GHL, FFH, and WZX revised the manuscript. All authors read and approved the final manuscript.

## Ethics declarations

### Ethics approval and consent to participate

Not applicable.

### Consent for publication

Not applicable.

